# Role of AAA-ATPase Cdc48p in Peroxisomal Quality Control

**DOI:** 10.1101/2024.10.14.618194

**Authors:** Ismaila Francis Yusuf, Wolfgang Girzalsky, Ralf Erdmann

## Abstract

The majority of peroxisomal matrix proteins are equipped with a type 1 peroxisomal targeting signal (PTS1), which is recognized in the cytosol by the import receptor Pex5p and targeted to the peroxisomal membrane. After cargo translocation into the peroxisomal lumen, the receptor is monoubiquitinated and recycled to the cytosol in an ATP-dependent manner by the AAA-ATPases Pex1p and Pex6p. Defects in receptor recycling trigger a quality control process by which the receptor is polyubiquitinated, extracted from the membrane and targeted to the proteasome for degradation by the RADAR (Receptor Accumulation and Degradation in the Absence of Recycling)-pathway.

Although the RADAR-pathway is conserved among species, it seemed to be missing in baker’s yeast. We present the identification and characterization of the RADAR pathway in *S. cerevisiae* and discover that the AAA-ATPases Msp1p and predominantly Cdc48p together with its co-factors Ufd1p/Npl4p are constituents of this pathway. In the RADAR-pathway, Cdc48p cooperates with the heterodimeric Ufd1p/Npl4p cofactor to pull misfolded, polyubiquitinated receptor out of the peroxisomal membrane for its subsequent degradation.

## Introduction

Peroxisomes are single membrane-bound organelles that are present in almost all eukaryotic cells. They carry out a wide range of metabolic functions depending on the organism and the cell type. A most conserved peroxisomal function is the degradation of fatty acids by β-oxidation and the detoxification of hydrogen peroxide (Smith & Aitchison, 2013, Wanders & Waterham, 2006). The dysfunction of peroxisome in humans or deficiency in the activity of peroxisomal enzymes leads to serious diseases that are often severe such as Zellweger syndrome and Rhizomelic chondrodysplasia punctata (RCDP) type 1) (Fujiki, Okumoto et al., 2022, Waterham & Ebberink, 2012). To maintain peroxisome homeostasis, molecular machines must import peroxisomal matrix proteins and membrane proteins encoded by nuclear genes and synthesized on free ribosomes in the cytosol (Lazarow & Fujiki, 1985). Peroxisomal matrix proteins are targeted by peroxisomal targeting signals, namely PTS1 or PTS2 (Kalel & Erdmann, 2018). The PTS1 is a tripeptide found at the C-terminus of the PTS1 cargo proteins and is recognized by the cycling receptor Pex5p. In contrast, the PTS2 is a nonapeptide located at the N-terminus of the corresponding PTS2 cargo protein and is recognized by the receptor Pex7p that works together with the co-receptors Pex18p or Pex21p in *S. cerevisiae* (Kunze, 2020). Subsequently to cargo recognition, The PTS receptor/cargo complex docks at the peroxisomal membrane with the docking complex consisting of Pex13p, Pex14p, and Pex17p (Albertini, Rehling et al., 1997, Huhse, Rehling et al., 1998). At the peroxisomal membrane, a dynamic and transient import pore is formed, which leads to the release of cargo proteins into the peroxisomal lumen (Erdmann & Schliebs, 2005). Upon the release of its cargo, the receptor is exported back to the cytosol to allow a new round of matrix protein import. This process depends on the monoubiquitination of the receptor at a conserved cysteine residue near its N-terminus (Platta, Brinkmeier et al., 2016) and is mediated by the E2 ubiquitin-conjugating enzyme Pex4p and the RING (really interesting new gene) E3 ligases Pex12p and Pex10p (Platta, El Magraoui et al., 2009, Platta, El Magraoui et al., 2007, Williams, van den Berg et al., 2007). The ATP-dependent export of the ubiquitinated receptor is facilitated by the peroxisomal AAA ATPase complex, Pex1p, and Pex6p (Miyata & Fujiki, 2005, Platta, Grunau et al., 2005). When recycling of Pex5p is blocked, e.g. as a result of a mutation in the exportomer components, yeast Pex5p is polyubiquitinated at a conserved Lys18 and Lys24 residue using the cytosolic E2 enzymes Ubc4p and the redundant Ubc5p and Ubc1p (Kiel, Emmrich et al., 2005, Platta, Girzalsky et al., 2004) as well as the E3-ligase Pex2p and Pex10p (El Magraoui, Baumer et al., 2012, Williams, van den Berg et al., 2008). The polyubiquitination of Pex5p and its extraction from the membrane has a quality control function by which dysfunctional and accumulated Pex5p is targeted for proteasomal degradation (Kiel et al., 2005). A block of Pex5p recycling leads to accumulation of the receptor at the peroxisomal membrane and triggers a quality control pathway known as RADAR (Receptor Accumulation and Degradation in the Absence of Recycling) by which the receptor is polyubiquitinated and degraded via the proteasome (Leon et al, 2006, Aksam et al., 2009, Leon & Subramani, 2007). However, the identity of the components of the RADAR pathway, especially proteins that extract polyubiquitinated Pex5p and mediates its degradation remained unknown.

Extraction of proteins from membranes in cells is performed by AAA-ATPases like Msp1 (ATAD1 in mammals) and Cdc48p (p97 or VCP in mammals). Msp1 is a membrane-anchored AAA-ATPase mainly localized to the outer mitochondria membrane and partially to peroxisomal membranes (Nakai, Endo et al., 1993, Wiese, Gronemeyer et al., 2007). Msp1p is required in protein quality control at the OMM, where it acts as a membrane dislocase for mistargeted tail-anchored protein (Chen, Umanah et al., 2014, Dederer & Lemberg, 2021, Okreglak & Walter, 2014). In addition, Msp1p has been shown to extract proteins that have become stuck in mitochondria import channels and target them for proteasomal degradation (Basch, Wagner et al., Weidberg & Amon, 2018). The function of peroxisome-associated Msp1p remains elusive. On the other hand, Cdc48p (p97/VCP in higher eukaryotes) is an essential AAA-ATPase involved in diverse cellular activities such as protein quality control (Saffert, Enenkel et al., 2017). Cdc48p extracts polyubiquitinated substrate polypeptides from membranes or macromolecule complexes in an ATP-dependent process and generally delivers them to the proteasome for degradation (Bodnar and Rapoport, 2017a; van den Boom and Meyer, 2018, Olszewski et al., 2019). The segregase activities of Cdc48p are required in diverse cellular processes such as ubiquitin fusion degradation (UFD) (Ghislain, Dohmen et al., 1996), endoplasmic reticulum-associated degradation (ERAD) (Hampton, 2002), mitochondria-associated degradation (MAD)(Heo, Livnat-Levanon et al., 2010), ribosome quality control (Verma, Oania et al., 2013), chromatin-associated degradation (CAD) (Dantuma & T., 2012), autophagy, and regulation of cell cycle progression (Böhm & A., 2013, Fu, Ng et al., 2003). Many cofactors regulate these different cellular pathways of Cdc48p by targeting it to specific substrate and subcellular localization (Buchberger, 2010, Buchberger, Schindelin et al., 2015).

Although the RADAR pathway is conserved among species, it seemed not to exist in baker’s yeast and its composition, e. g. the machinery that extracts non-recycled or accumulated and polyubiquitinated Pex5p from the peroxisomal membrane remained unknown. Here, we present the identification and characterization of the RADAR pathway in *S. cerevisiae* and show that the AAA-ATPases Msp1 as well as Cdc48p and its cofactors Ufd1p/Npl4p are major components of this pathway that is required to extract dysfunctional Pex5p from the peroxisomal membrane for its proteasomal degradation. Accordingly, Cdc48p cooperates with the heterodimeric Ufd1p/Npl4p cofactor to pull misfolded, polyubiquitinated receptor out of the peroxisomal membrane for their subsequent degradation.

## Results

### Pex5p is rapidly degraded in cells lacking *PEX1* during the exponential growth phase

Upon peroxisomal protein import, the PTS1 receptor Pex5p is monoubiquitinated and recycled after cargo translocation into the lumen of peroxisome (El Magraoui, Brinkmeier et al., 2013, Platta et al., 2005). In this scenario the release of the monoubiquitinated receptor is performed by the AAA-ATPases Pex1p and Pex6p, which form a heterohexameric complex and are bound to the peroxisomal membrane by Pex15. A block of Pex5p recycling leads to accumulation of the receptor at the peroxisomal membrane and triggers a quality control pathway known as RADAR (Receptor Accumulation and Degradation in the Absence of Recycling) by which the receptor is polyubiquitinated and degraded via the proteasome (Leon et al., 2006, Aksam et al., 2009, Leon & Subramani, 2007). Accordingly, based on steady-state protein levels, Pex5p was reported to undergo rapid degradation in human cells and several yeast strains, including *P. pastoris*, in the absence of Pex1p/Pex6p, with the consequence of a significantly decreased Pex5p-level, when compared to wild type cells (Collins, Kalish et al., 2000, Yahraus et al., 1996, Leon et al, 2006). In contrast, under the same conditions, a wild type like level of Pex5p was detected in the corresponding mutants of *S. cerevisiae* (Kiel et al., 2005; Platta et al., 2004). In these species, poly-ubiquitinated Pex5p accumulated at the peroxisomal membrane. We were puzzled by this observation and the conclusion that the RADAR-pathway might not be active in *S. cerevisiae.* We considered that the differences in Pex5p turnover of baker’s yeast might be due to different growth conditions. To this end, we compared the steady concentration of Pex5p from cells grown to stationary phase with those of the exponential growth phase. Wild type yeast cells were compared with cells lacking the receptor export component Pex1p (*pex1Δ*), or a component of docking complex (*pex17Δ*) or the PTS1-receptor (*pex5Δ*). Pex5p-level were estimated by immunoblotting, while detection of mitochondrial porin served as loading control (**Figure 1A/1B**). Protein levels were quantified by signal intensity measurements (**Figure 1C**). The Pex5p-steady-state level of wild type and *pex1Δ* cells was similar under both conditions, while that of *pex17Δ* cells showed an increased level (**Figure 1**). This result indeed indicated that the RADAR pathway may not be active in *S. cerevisiae*. However, as the steady-state level of a protein is a result of its expression rate related to the speed of degradation, we performed additional experiments that aimed to analyze the degradation process separately. To this end, we used cycloheximide (CHX), an antibiotic that inhibit protein biosynthesis by blocking translational elongation at the ribosome (Obrig et al., 1971). To perform the cycloheximide-chase, the antibiotic was added to the growth medium of oleic acid-induced cells that were grown to stationary (OD600nm = 2.5) or exponential phase (OD600nm =1.5). Cells incubated with dimethylsulfoxid (DMSO) served as a control and samples were taken at different time points as indicated (**Figure 1D/1E**). The analysis of Pex18p-degradation served as a control for the block of protein translation, as it permanently undergoes rapid degradation (Purdue & Lazarow, 2001). In line with (El Magraoui et al., 2013, Purdue & Lazarow, 2001), Pex18p was degraded in wild type as well as in *pex1Δ* cells. Pex18p was degraded in these cells from the exponential as well as from the stationary growth phase (**Figure 1D/1E**). In contrast, Pex5p remained stable in *pex1Δ* cells grown at stationary growth phase (**Figure 2D**), which is in line with reported findings for *S. cerevisiae* (Kiel et al., 2005, Platta et al., 2004). Remarkably, rapid degradation of Pex5p was visible in cells of the exponential growth phase (**Figure 2E**). The quantification of Pex5p signal intensity revealed a reduction of Pex5p of about 80% when CHX was added. This result showed that the degradation rate of *S. cerevisiae* Pex5p depends on the growth conditions and indicated that also *S. cerevisiae* does possess the RADAR pathway, which therefore is conserved among species.

**Figure 1:**
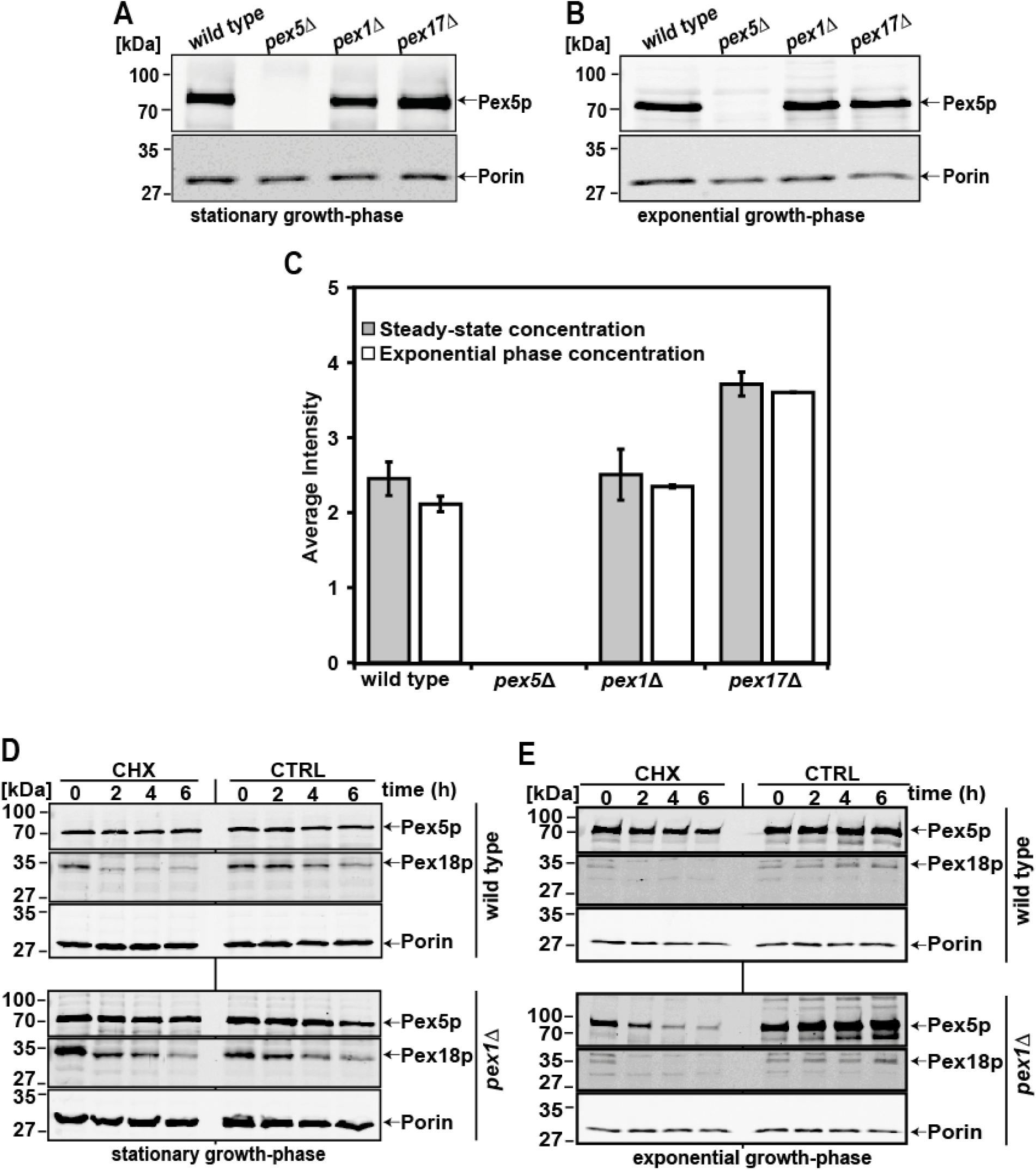
Pex5p degradation in cells lacking *PEX1* is dependent on the growth phase of *S. cerevisiae*. Steady-state level of Pex5p in wild type, *pex5Δ*, *pex1Δ* and *pex17Δ* cells in different growth phases were analysed by immunoblotting using Pex5p-specific antibodies (A-C). Cell lysates were derived from cells grown in oleic acid media to stationary phase (A), or exponential phase (B). Mitochondrial porin served as loading control. Wild type and *pex1Δ* cells show a similar Pex5p steady-state concentration at both stationary and exponential growth phases, while the amount of Pex5p in *pex17Δ* cells is slightly increased (C). The intensities of the Pex5p signal of three independent experiments were quantified densitometrically, normalized to porin intensities and graphically displayed. Turnover of Pex5p in different growth phases (D and E). Wild type and *pex1Δ* cells were grown in oleic acid-containing media to stationary phase (OD_600nm_ = 2.0-2.5) (D) or exponential phase (OD_600nm_ = 1-1.5) (E) and then treated with DMSO (Ctrl) or 0.5 mg/ml cycloheximide (CHX). Cells lysates were prepared by TCA precipitation at the indicated times and proteins were analyzed by immunoblotting using the indicated antibodies. The graph below shows Pex5p signal intensities normalized to porin and plotted against the chase time with Pex5p protein amount at T0 set to 1. At stationary phase, Pex5p is relatively stabilized in cells lacking *PEX1*. In contrast, Pex5p is rapidly degraded in the exponential growth phase. Error bars = S.E.M with n = 3

**Figure 2:**
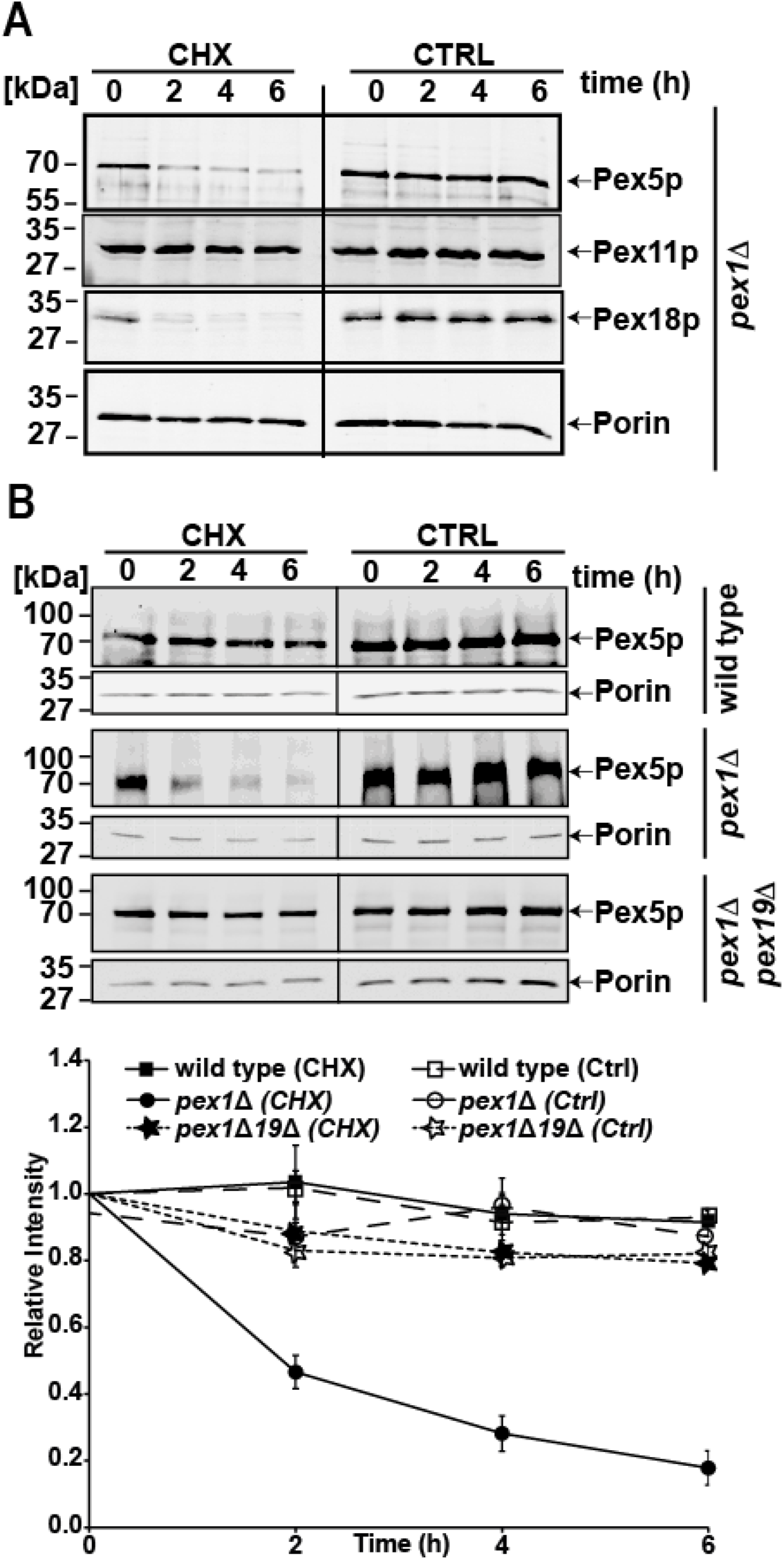
Pex5p degradation in cells lacking *PEX1* is not an effect of pexophagy and requires the peroxisomal membrane. *S. cerevisiae* cells were grown in oleic acid-containing media to exponential phase (OD_600nm_ = 1-1.5) and then treated with DMSO (Ctrl) or 0.5 mg/ml cycloheximide (CHX). Cell lysates were prepared by TCA precipitation at the indicated times and proteins were analyzed by immunoblotting using indicated antibodies. (A) In *pex1Δ* cells, the stability of the peroxisomal membrane marker Pex11p in comparison to Pex5p was analysed. Here, the amount of Pex11p remained unchanged, indicating that that the decrease in Pex5p with time is not due to pexophagy. (B) Analyses of Pex5p protein amount in *pex1Δpex19Δ* cells relative to WT and *pex1Δ* cells. The graph below represents the Pex5p signal intensities, which were normalized to porin and plotted against the chase time. The Pex5p protein amount at T0 was set to 1. The observed stability of Pex5p in *pex1Δpex19Δ* knockout cells proves that peroxisomal membrane is required for Pex5p turnover. Error bars = S.E.M with n = 3.

### Degradation of Pex5p is not due to pexophagy and requires the peroxisomal membrane

Inspired by the finding that Pex5p is actively degraded in *S. cerevisiae pex1*Δ cells, we sought to identify factors that contribute to Pex5p turnover. It has been shown that the loss of peroxisomal AAA-ATPase complex could lead to an increased peroxisome breakdown (Law, Bronte-Tinkew et al., 2017, Nuttall, Motley et al., 2014). This autophagic degradation of entire peroxisomes termed as pexophagy is indicated by a simultaneous degradation of peroxisomal matrix and membrane proteins. In yeast, pexophagy can be induced by nitrogen starvation (Klionsky, Abdelmohsen et al., 2016) and degradation of the peroxisomal membrane protein Pex11p can function as a measure for this process (Mastalski, Brinkmeier et al., 2020). To exclude that the observed Pex5p degradation in the RADAR pathway is caused by pexophagy, *pex1*Δ cells were grown in presence of nitrogen in exponential growth rate in presence and absence of CHX and whole cell lysates were prepared at indicated time points (**Figure 2A**). As shown before (**Figure 1E**), both Pex5p as well as Pex18p display decreased steady-state concentrations when CHX was added to the growth media, whereas both proteins remained stable without CHX. Pex11p was stable under both conditions, demonstrating that under our chosen conditions, no pexophagy was induced, and Pex5p was degraded by another process (**Figure 2A**).

To provide the platform for the identification of proteins that could be required for Pex5p turnover, we investigated the influence of the peroxisomal membrane on Pex5p turnover. Pex5p export components such as the AAA-complex are membrane-localized and we reasoned that polyubiquitinated Pex5p that is destined for proteasomal degradation must be extracted from the membrane and delivered to the proteasome, which is mostly cytosolic (Platta et al., 2005, 2003). To elucidate the influence of the peroxisomal membrane, we monitored Pex5p turnover in *pex1Δpex19Δ* double knockout cells. Pex19p per se is the receptor and/or chaperone for peroxisomal membrane proteins, required for peroxisomal membrane biogenesis (Jansen & van der Klei). Cells lacking Pex19p are characterized by the lack of peroxisomes or peroxisomal remnants so called ghosts (Koek, Komori et al., 2007). For this reason, Pex5p is unable to functionally dock to the peroxisomal membrane via the receptor docking complex, composed of Pex13p, Pex14p and Pex17p (Rüttermann & Gatsogiannis, 2022). CHX chase experiments of *pex1Δpex19Δ* double knockout cells revealed that Pex5p concentration remained stable, as seen in wildtype cells (**Figure 2B**). In contrast, signal intensity measurement revealed a degradation of about 80 % of Pex5p in *pex1Δ* cells in the presence of CHX. This result showed that the absence of Pex19p and thus the absence of peroxisomal membranes results in a stabilization of Pex5p. It therefore can be concluded that Pex5p attachment to the peroxisome membrane is a prerequisite for a later handover to the still unknown degradation machinery of the RADAR-pathway.

### The AAA-ATPase Msp1p has an influence on Pex5p turnover

Next, we sought to identify proteins required to extract Pex5p from the membrane and facilitate its proteasomal degradation in cells lacking Pex1p. In this context we focused on AAA-proteins that have been shown to extract proteins from their native environment and mediate their proteasomal turnover. This included the mitochondrial sorting protein Msp1 (ATAD1 in mammals). Msp1p is localized to the mitochondria’s outer membrane (OMM) and partially to the peroxisomal membrane (Nakai et al., 1993, Wiese et al., 2007). Required in protein quality control at the OMM, Msp1p acts as a membrane dislocase for mistargeted tail-anchored proteins (Chen et al., 2014, Okreglak & Walter, 2014). In addition, Msp1p has been shown to extract proteins that stuck in mitochondria import channels and target them for proteasomal degradation (Weidberg & Amon, 2018). Here we investigated whether Msp1p facilitates Pex5p turnover in the absence of Pex1p. Based on our CHX chase experiments, we observed that in strains lacking Msp1p alone, no effect on Pex5p protein amount was visible, since the level of Pex5p in *msp1Δ* cells is comparable to that of wild type (**Figure 3**). The absence of Pex1p did result in the established degradation of Pex5p. However, the addition knockout of Msp1p caused a partial stabilization of Pex5p, which was confirmed by quantification of Pex5p signal intensity (**Figure 3, right panel**). This result indicates that Msp1p may play a role in Pex5p quality control as the Pex5p amount of *pex1Δmsp1Δ* cells was slightly higher than those of *pex1Δ* cells. However, the data also leave no doubt that the absence of Msp1 did hamper the degradation of Pex5p only slightly. From this result we conclude that other components than Msp1 are responsible for the turnover of the bulk of Pex5p.

**Figure 3:**
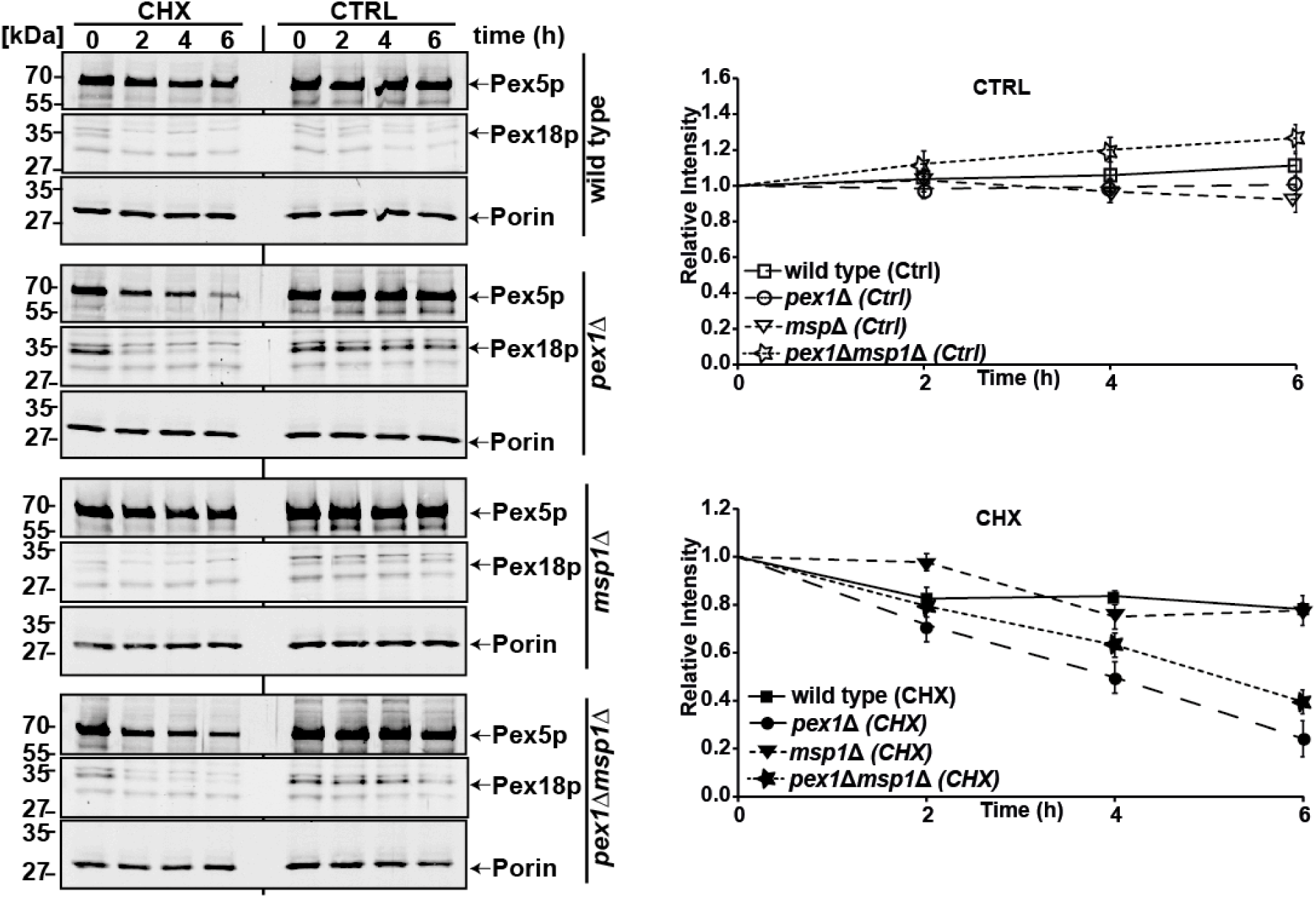
Role of Msp1p on Pex5p degradation in the RADAR pathway. Indicated strains were grown in oleic acid-containing media to exponential phase (OD_600nm_ = 1-1.5) and then treated with DMSO (Ctrl) or 0.5 mg/ml cycloheximide (CHX). Cell lysates were prepared by TCA precipitation at the indicated times and proteins were analyzed by immunoblotting using indicated antibodies. The graph on the right represents the Pex5p signal intensities, which were normalized to porin and plotted against the chase time. Pex5p protein amount at T0 was set to 1. As observed for *msp1Δpex1Δ* double knockout cells, Pex5p protein amount is slightly increased compared to *pex1Δ* cells, suggesting that Msp1 might have a partial effect on Pex5p turnover. Error bars = S.E.M with n = 3.

### Construction of Cdc48p conditional knockout strains

As Pex5p is still degraded in the absence of Msp1p, our data are clear that another factor must be involved in the degradation process. A likely candidate was Cdc48p (p97/VCP in higher eukaryotes), which is an essential AAA-ATPase involved in diverse cellular activities such as protein quality control (Saffert et al., 2017). Cdc48p/VCP is involved in various cellular processes like protein degradation, membrane fusion and chaperone activity. Increased levels of Cdc48p/VCP correlate with cancer, whereas Cdc48p/VCP at endogenous levels has been proposed to be a pathological effector in protein deposition diseases. In the past, it turned out that yeast is a suitable model organism to analyze Cdc48p functions. By now it is clear that Cdc48p recognizes polyubiquitinylated proteins and facilitates their dissociation from protein complexes and organelle membranes using the energy derived from ATP hydrolysis (Hanson & Whiteheart, 2005). The segregase activity of Cdc48p is required for diverse cellular processes such as ubiquitin fusion degradation (UFD) (Ghislain et al., 1996), endoplasmic reticulum-associated degradation (ERAD) (Hampton, 2002), mitochondria-associated degradation (MAD) (Heo et al., 2010), ribosome quality control (Verma et al., 2013), chromatin-associated degradation (CAD) (Dantuma & T., 2012), autophagy, and regulation of cell cycle progression (Böhm & A., 2013, Fu et al., 2003). Many cofactors regulate these different cellular pathways of Cdc48p by organizing its targeting to specific substrates and subcellular localizations (Buchberger, 2010, Hänzelmann & Schindelin, 2017).

However, the work with Cdc48p is hampered by the fact that this protein is essential. Thus, studies in yeast were performed with conditional mutants, mostly temperature sensitive (ts) mutants. However, sometime these ts-mutants are difficult to work with as the proteins may not be fully functional even at the permissive temperature or other stress conditions like growth on oleic acid. To address a possible peroxisomal function of the essential Cdc48p, we therefore generated a conditional knock-out strain. To this end, we genomically integrated the tightly regulated GAL1-promoter in front of the CDC48 open reading frame (**Supplementary Figure 1**), which allowed to switch off the expression of the essential CDC48.

To confirm proper expression-regulation of CDC48 via the Gal1-promotor, cells of the wild type and the newly generated strain (GAL1-CDC48) were first grown on glucose-medium, subsequently shifted to either glucose or galactose-containing medium of different concentrations and whole cell lysates were analyzed for Cdc48p-abundance. Mitochondrial porin served as an internal loading control and did not differ between the analyzed samples (**Supplementary Figure 2**). In wild type, the Cdc48p level was nearly the same under the changing glucose concentrations, whereas no Cdc48p sensitive signal was obtained in samples of the GAL1-CDC48 even at low glucose conditions. In addition, cells of GAL1-CDC48 showed a significant growth defect in glucose media as expected for a strain lacking an essential protein (**Supplementary Figure 2A**). When GAL1-CDC48 cells were cultivated under different concentration of galactose, an induction of Cdc48p expression was observed, which increased with increasing galactose in the medium. Under these conditions, the GAL1-CDC48 strain displayed a wild type like growth rate (**Supplementary Figure 2B**). We also tested Cdc48p expression and growth behavior of the strain in media containing oleate alone or supplemented with either glucose or galactose (**Supplementary Figure 2C**). The results showed that the expression of Cdc48p is only induced in the presence galactose, which is also supported by its growth behavior in galactose containing media. All together, these results clearly demonstrated a tight glucose repression and efficient galactose-induction of Cdc48p in the GAL1-CDC48 conditional knockout strain.

### The AAA-ATPase Cdc48p is required for Pex5p quality control

The newly generated GAL1-CDC48 conditional knockout strain was applied to analyze the impact of Cdc48p in Pex5p quality control. To this end, we monitored the levels of Pex5p in RADAR-induced cells in the presence and absence of Cdc48p. Pex18p served as a control for the block of protein translation since it is rapidly degraded even under wild type conditions (Hensel, Beck et al., 2011). As knockdown of an essential gene like CDC48 could cause several pleiotropic effects that could affect cell viability, we also checked Fox3p (thiolase) levels (**Figure 4**). Fox3p is highly induced by oleate and its increase over time illustrated the proper uptake of this fatty acid and indicated that the cells are still healthy enough for protein synthesis even upon depletion of Cdc48p for 8 hours. As outlined above, the peroxisomal membrane protein Pex11p was analyzed as a measure for pexophagy. Mitochondrial porin was used as loading control.

**Figure 4:**
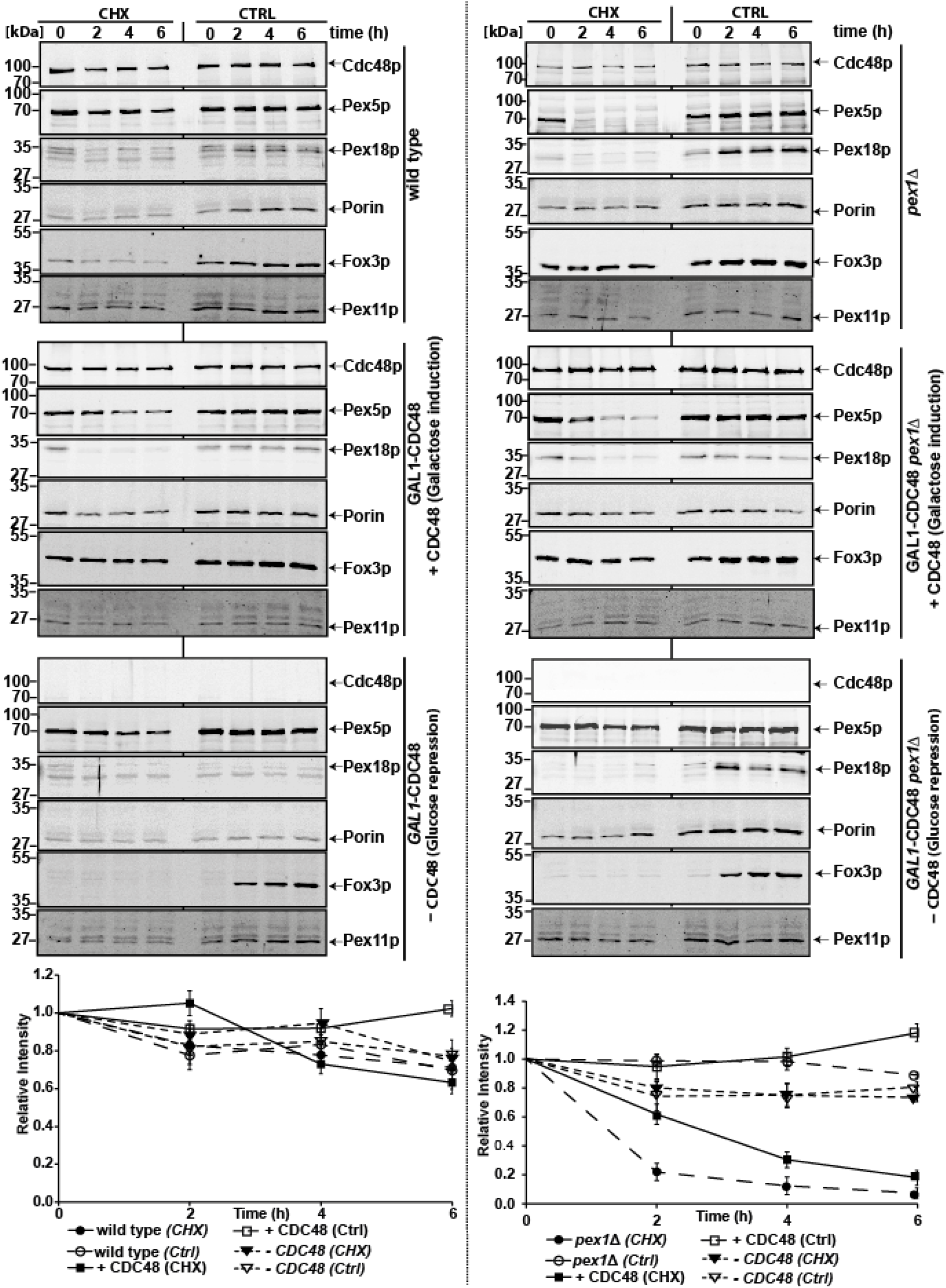
Role of Cdc48p for Pex5p degradation in the RADAR pathway. Wildtype cells and *pex1Δ*, and corresponding GAL1-CDC48 integrated cells were cultivated in media containing 0.1 % glucose and 0.1 % oleate (repressing condition) or 0.1% oleate and 0.1 % galactose to induce Cdc48p expression. All strains were cultivated to the exponential phase (OD600nm = 1-1.5) and then treated with DMSO (Ctrl) or 0.5 mg/ml cycloheximide (CHX). Cell lysates were prepared by TCA precipitation at indicated times, and proteins were analysed by immunoblotting using indicated antibodies. The graph below each panel shows Pex5p signal intensities normalised to porin and plotted against the chase time. Pex5p protein amount at T0 was set to 1. Induction or knockdown of Cdc48p in the wild type background had no effect on Pex5p protein amount. However, in GAL1-CDC48 *pex1Δ* cells under inducing CDC48 expression, Cdc48p was present and Pex5p was rapidly degraded, while block of expression and depletion of Cdc48p inhibited Pex5p degradation. Error bars = S.E.M with n = 3.

In wild type cells, the genomic expression of Cdc48p had no effect on the Pex5p protein level over time when compared to native wild type cells (**Figure 4, left panel**). In both cases only a slight decrease of Pex5p was observed with time.

As described above, deletion of Pex1p resulted in a rapid degradation of Pex5p (**Figure 4, right panel, *pex1Δ***). Equally, induction of Cdc48p-expression in *pex1Δ* cells led to Pex5p degradation as seen for *pex1Δ* cells (**Figure 4, right panel, *pex1Δ* +CDC48**). Thus, in cells lacking Pex1p, Pex5p is rapidly degraded in the presence of Cdc48p. Interestingly, repression of Cdc48p expression in *pex1Δ* cells reflecting in principle a *pex1Δcdc48Δ* double-deletion led to significant impaired Pex5p degradation (**Figure 4, right panel, *pex1Δ* -CDC48**). Thus, in the absence of Cdc48, the degradation of Pex5p is blocked. Turn-over of Pex18p illustrated that protein degradation per se was functional even in absence of Cdc48p. Pex11p stability excludes enhanced pexophagy in absence of Cdc48p (**Figure 4, left panel)**.

Quantification revealed that in the presence of Cdc48p in the *pex1Δ*-mutant, about 90% of Pex5p was degraded, while in the absence of Cdc48p less than 20% of Pex5p was degraded (**Figure 4, right lower panel**). The block in export also did result in an accumulation of poly-ubiquitinated Pex5p (**Supplementary Figure 3**). These data put Cdc48p in the leading role as being responsible for the Pex5p degradation in the RADAR pathway, with Msp1p probably playing only a minor role. To investigate this in more detail, we extended our analysis of the absence and presence in the double knockout strain of *pex1Δmsp1Δ*-GAL1-CDC48. We observed that in the presence of Cdc48p in the double knock-out strain *pex1Δmsp1*, most of Pex5p is still degraded in the presence of CHX, while upon expression of Cdc48p, the degradation of Pex5p is severely blocked. (**Figure 5**). The quantification revealed that the additional depletion of Msp1 increased stabilization to 85% (**Figure 5, lower panel**), which is slightly more than the stabilization of about 80% of Pex5p in *pex1Δ cells* upon Cdc48p depletion (**Figure 4, right lower panel**). However, the strong stabilization of Pex5p upon depletion of Msp1 and Cdc48p suggests that no other AAA-ATPase seems to be involved in this process. Moreover, the data indicate that Cdc48p plays the major role in Pex5p degradation in the RADAR pathway, with a redundant and slight impact of Msp1p.

**Figure 5:**
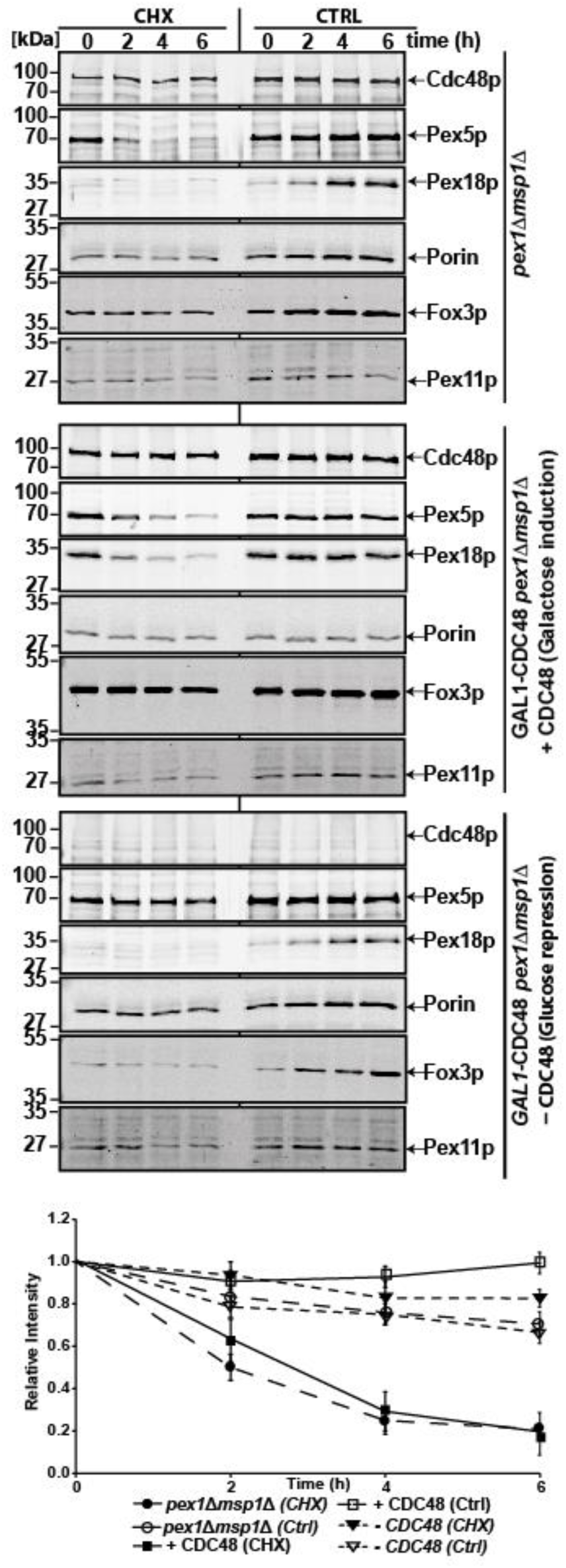
Cdc48p is the major AAA-ATPase in the RADAR pathway. *pex1Δmsp1Δ* cells (i) were cultivated in media containing 0.1 % glucose and 0.1 % oleate. GAL1-CDC48 *pex1Δmsp1Δ* cells were cultivated in media containing 0.1% oleate and 0.1 % galactose to induce Cdc48p expression or cultivated in media containing 0.1% oleate and 0.1 % glucose to deplete Cdc48p. All strains were cultivated to the exponential phase (OD600nm = 1-1.5) and then treated with DMSO (Ctrl) or 0.5 mg/ml cycloheximide (CHX). Cell lysates were prepared by TCA precipitation at the indicated times, and proteins were analysed by immunoblotting using indicated antibodies. The graph below shows Pex5p signal intensities normalised to porin and plotted against the chase time after the Pex5p protein amount at T0 was set to 1. The data show that in the absence of both AAA-proteins, Pex5p is not degraded, indicating that no other AAA-ATPase seems to be involved in the RADAR pathway. In the absence of Msp1 but presence of Cdc48p, Pex5p is rapidly degraded, indicating that Cdc48p plays a major role in Pex5p degradation. Error bars = S.E.M with n = 3.

### The peroxisomal RADAR-pathway depends on Cdc48p co-factors Npl4p and Ufd1p

Cdc48p is recruited to substrates and targeted to distinct cellular sites by different cofactor proteins that mediate its ubiquitin and proteasome binding (Jentsch & Rumpf, 2007). Npl4p and Ufd1p are Cdc48p cofactors required for the degradation of Cdc48p substrates in endoplasmic reticulum and mitochondria (Buchberger, 2010, Metzger, Scales et al., 2020). Due to the functional connection of peroxisome to both cellular compartments, we examined whether these Cdc48p co-factors are required for Pex5p turnover in the RADAR-pathway.

Like Cdc48p, Npl4p and Ufd1p are both essential for cell survival. To test their involvement in Pex5p turnover, we followed the same strategy that was used for knockdown of Cdc48p. Thus, conditional knockout strains of Npl4p and Ufd1p were generated by the genomic integration of the galactose promoter upstream of these genes to control their expression. The observed expression of oleate-inducible Fox3p served as a proof of cell viability, which is expected to be affected by the knockdown of these essential genes. Based on CHX chase experiments, knockdown of Npl4p as well as Ufd1p did result in a stabilization of Pex5p a *pex1Δ* background (**Figure 6**). In addition, polyubiquitinated Pex5p accumulated in Npl4p knockdown cells (**Supplementary Figure 3**). The data indicate that the Npl4p-Ufd1p complex mediates the function of Cdc48p in Pex5p turnover. UBX family of proteins have been implicated in Cdc48p function, where they have been shown to recruit the Cdc48p-Npl4p-Ufd1p complex to substrate. For this reason, we tested whether Ubx2p, Ubx3p, Ubx5p and Vms1p, which has been implicated in protein quality control, are required for Pex5p degradation. However, the deletion of these *UBX* genes and *VMS1* did not affect Pex5p degradation (**Supplementary Figure 4**), indicating that these proteins are not required for Pex5p turnover.

**Figure 6:**
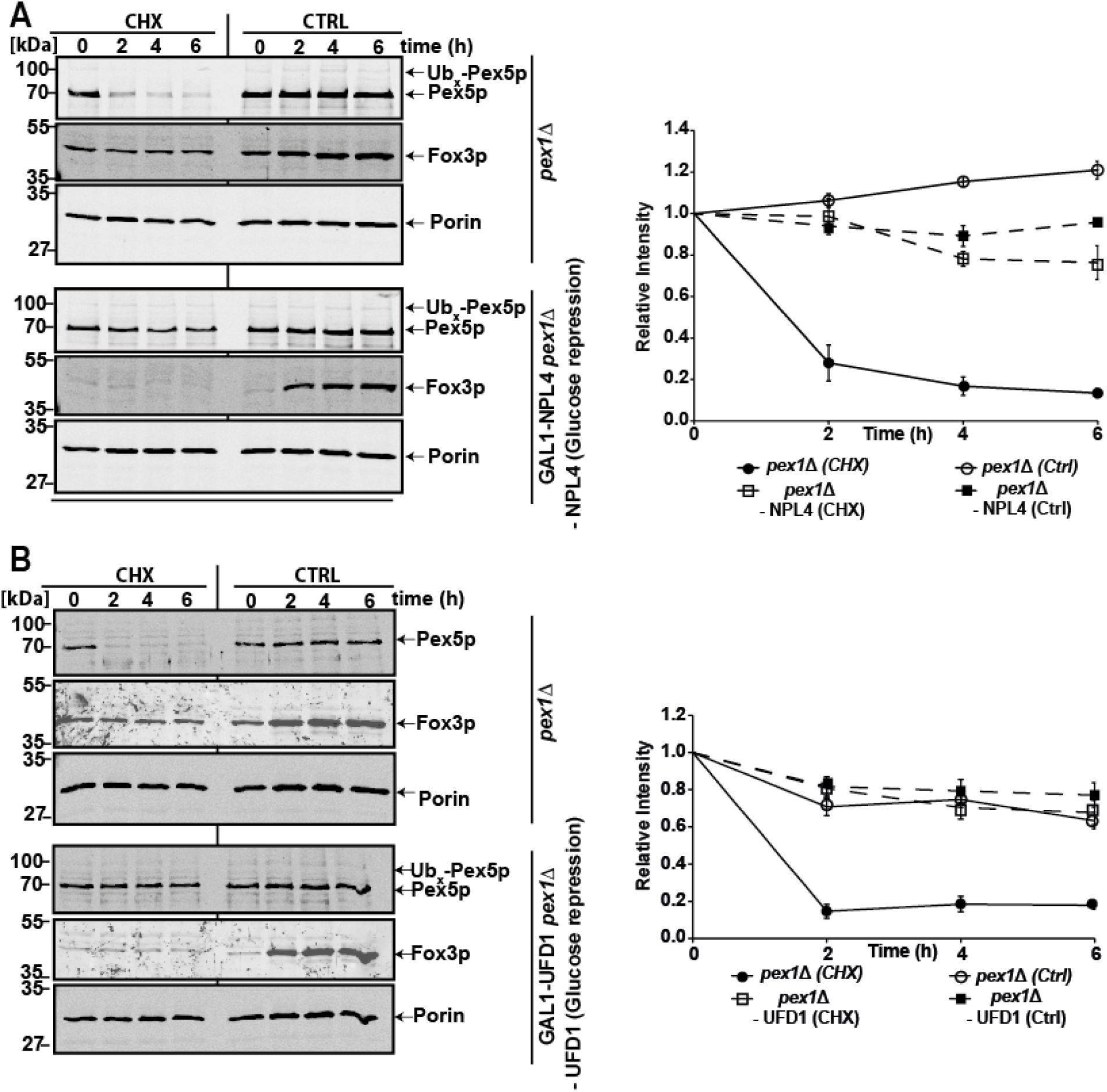
The Cdc48p co-factors Npl4p and Ufd1p are required for Pex5p quality control. Mutant cells of *pex1Δ*, GAL1-NPL4 *pex1Δ* and GAL1-UFD1 *pex1Δ* were cultivated in media containing 0.1 % glucose and 0.1 % oleate to deplete Npl4p (A) or Ufd1p (B). All strains were cultivated to the exponential phase (OD600nm = 1-1.5) and then treated with DMSO (Ctrl) or 0.5 mg/ml cycloheximide (CHX). Cell lysates were prepared by TCA precipitation at the indicated times, and proteins were analysed by immunoblotting using the indicated antibodies. The graph beside each blot shows Pex5p signal intensities normalised to porin and plotted against the chase time after the Pex5p protein amount at T0 was set to 1. Error bars = S.E.M with n = 3.

## Discussion

Pex5p is a crucial cycling receptor involved in the import of peroxisomal matrix proteins and plays a pivotal role in the biogenesis of peroxisomes. Through its interaction with peroxisomal targeting signal type 1 (PTS1) proteins, Pex5p mediates their translocation into the peroxisomal matrix. In all organisms, Pex5p undergoes a dynamic cycle of import from the cytosol to the peroxisome and export back to the cytosol. The import process involves recognition of PTS1 cargo proteins and subsequent docking onto the peroxisomal membrane for translocation. After cargo delivery, Pex5p is ubiquitinated, initiating recycling back to the cytosol. In addition to the ubiquitination machinery, the export-processes requires the AAA-proteins Pex1p/Pex6p (Platta, Hagen et al., 2014). Dysregulation of Pex5p cycling due to a defect in the receptor export machinery triggers a degradation or quality control pathway known as RADAR (Receptor Accumulation and Degradation in Absence of Recycling) that is required in regulating Pex5p turnover in peroxisome biogenesis (Leon et al, 2006; Aksam et al., 2009, Leon & Subramani, 2007). Although the RADAR pathway is conserved among species, it seemed not to exist in baker’s yeast and its composition, e.g. the machinery that extracts non-recycled or accumulated and polyubiquitinated Pex5p from the peroxisomal membrane remained unknown. Here, we present the identification and characterization of the RADAR pathway in *S. cerevisiae* and show that the AAA-ATPase Cdc48p and its cofactors Ufd1p/Npl4p are major components of this pathway that is required to extract dysfunctional Pex5p from the peroxisomal membrane for its proteasomal degradation.

### Pex5p-undergoes rapid degradation in cells affected in the Pex1p/Pex6p-complex

A defect in receptor recycling in *P. pastoris* and human cells leads to drastically reduced steady state levels of Pex5p (Collins et al., 2000, Yahraus et al., 1996), which was explained by the RADAR-pathway (Leon et al, 2006) that is responsible for removal and proteasomal degradation of dysfunctional Pex5p. On first sight, it seemed that this pathway does not exist in baker’s yeast. Instead, Pex5p was shown to be polyubiquitinated followed by its accumulation at the peroxisomal membrane, which allowed the discovery of receptor poly-ubiquitination upon a defect in recycling (Kiel et al., 2005, Platta et al., 2004). Here we discovered that the RADAR pathway exists in baker’s yeast. It is active in cells grow at the exponential phase, but inactive in stationary phase cells, which does explain why this pathway has not been discovered earlier. This raises the question why disposal of dysfunctional Pex5p is limited to the exponential growth phase. A possible explanation could be related to the high level of metabolic activity during this phase and the corresponding steady demand for a functional peroxisomal protein import machinery, especially under oleic acid induction conditions. Thus, in exponentially grown cells, the maintenance of peroxisomal function might be a primary goal, and for this purpose the RADAR-pathway clears the import machinery of dysfunctional receptors. Accordingly, non-recycled Pex5p is actively targeted for degradation as the cells attempt to prevent the accumulation of non-functional Pex5p to maintain peroxisome function. Under this condition also pexophagy is repressed. It was shown that latent activation of the yeast pexophagy receptorAtg36 by the casein kinase Hrr25 in rich media is repressed by the ATPase activity of Pex1p/Pex6p (Yu et al, 2022), and yeast peroxisomes are constitutively degraded by selective autophagy in the absence of receptor recycling (Nuttall et al., 2014). However, this degradation is triggered by starvation, which explains that it is not evident in exponentially grown cells.

In the stationary growth phase, limiting resources cause a slow-down of cellular metabolism. In this situation, a major goal is the regeneration of resources, while the maintenance of peroxisome function might be secondary. This is expected to lead to an inhibition of the RADAR-pathway and the resulting lack of degradation is supposed to be responsible for the observed accumulation of polyubiquitinated Pex5p at the peroxisomal membrane (Platta et al., 2004, Kiel et al., 2005)

### Role of Msp1 in Pex5p degradation

Our study demonstrates that Cdc48p, Ufd1p, Npl4p and Msp1 are required for Pex5p degradation in the RADAR pathway (**Figures 5, 6**). Msp1p is a AAA-protein that is dually localized to the mitochondrial outer membrane and the peroxisomal membrane. Msp1 is conserved from yeast to humans and its orthologue ATAD1 has been shown to remove mislocalized or misfolded tail-anchored proteins from the mitochondrial outer membrane, such as mislocalized Pex15p in *S. cerevisiae* or Pex26 in humans (Chen et al., 2014, Okreglak et al., 2014, Nakai et al., 1993, Wang et al., 2022). As Msp1 is responsible for the removal of integral membrane proteins, its involvement in the degradation of the import receptor Pex5p might come as a surprise. However, it has been demonstrated that the accumulated Pex5p in *pex1Δ* cells behaves like an integral membrane protein and might be a target of Msp1p (Platta et a., 2005). In fact, it has recently been shown that ATAD1 is involved in Pex5p extraction from the peroxisomal membrane in human cells (Ott et al., 2022). However, our data are clear in that Pex5p is still rapidly degraded in strains lacking both Pex1p and Msp1p, which indicates that Msp1 is not the key-component of this quality pathway.

### Cdc48p-Ufd1p/Npl4p is required for Pex5p degradation

Cdc48p, Ufd1p and Npl4p function in concert in the degradation of dysfunctional ER proteins in the ERAD pathway. As these proteins are essential for cell viability, we used conditional knockout cells, which express the corresponding genes under the tight control of the GAL1 promoter. The knockdown of either CDC48, UFD1 or NPL4 abolishes Pex5p degradation, thereby maintaining Pex5p protein amount at wild type level. This result raised the question of whether the observed degradation is due to the absence of components that are involved in Pex5p degradation or if the cells are generally non-functional. Several observations show that cellular functions are preserved during the six hours of depletion. First, oleic acid induction and the corresponding increasing expression of Fox3p still takes place even upon depletion of the proteins. Moreover, degradation of Pex18p, which functions independently of the RADAR-pathway is not affected. Finally, the chosen growth conditions (0.1% glucose, 0.1% oleic acid) did result in a depletion of Cdc48p below the detection limit, but the cells did still grow for some hours, although very slowly (**Supplementary Figure 2**).

Cdc48p binds Ufd1 and Npl4 and this complex extracts ubiquitinated proteins from the ER membrane and facilitates their degradation via the cytosolic proteasome (Buchberger, 2010, Vembar & Brodsky, 2008, Ye, Meyer et al., 2003). In analogy, it can be expected that Cdc48p might function in the same way in the RADAR-pathway. However, it has been shown that Msp1 extracts mistargeted tail-anchored (TA) proteins from the mitochondrial outer membrane for reinsertion into the ER membrane. The ER-membrane-inserted TA proteins are then subjected to the ER quality control system, i.e., ubiquitination by Doa10 and subsequent re-extraction by Cdc48p to the cytosol for proteasomal degradation (Matsumoto et al, 2019). However, the ubiquitin ligases Pex2p, Pex10p and Pex12p that are responsible for the ubiquitination of Pex5p form a stable complex that is located at the peroxisomal membrane ((Tan et al, 1995; Kalish et al, 1995; Waterham et al, 1996; Kalish et al, 1996; El Magraoui et al, 2012; Feng et al, 2022). Moreover, poly-ubiquitinated Pex5p has been shown to accumulate at the peroxisomal membrane (Law et al., 2017), indicating that the place of ubiquitination of Pex5p is at the peroxisomal membrane and that Pex5p is not routed to the ER for degradation.

Cdc48p can interact with many cofactors that regulate its function by recruiting it to different cellular pathways. So far, about 40 cofactors have been identified in mammals, and the number is still growing (Buchberger, 2010). Our study show that the Cdc48p cofactors Ufd1p and Npl4p are also involved in Pex5p degradation. (**Figure 6**). Based on these results, we conclude that the Cdc48pUfd1/Npl4p complex is recruited to the peroxisomal membrane in cells lacking PEX1 to extract accumulated Pex5p and facilitate its proteasomal degradation in a way similar to ERAD. However, the diverse cellular functions of Cdc48p are often regulated by additional cofactors. In ERAD, UBX cofactors mediate the interaction between Cdc48p and ubiquitinated substrates (Buchberger, 2010). For instance, Ubx2 recruits the Cdc48pUfd1/Npl4p complex to the ER membrane to extract ubiquitinated proteins that are to be degraded by the proteasome (Neuber, Jarosch et al., 2005). Along this line, we checked whether the Cdc48p adaptors Ubx2p, Ubx3p, Ubx5p and Vms1p are required to degrade Pex5p. However, none of these adaptors turned out to be required for the RADAR-pathway.

The RADAR pathway was also suggested to target the PTS2 co-receptors Pex18p and Pex20p for proteasomal degradation (Purdue & Lazarow, 2001). However, this study shows that Cdc48p is not required for the degradation of *S. cerevisiae* Pex18p, suggesting that an alternative pathway exists for the degradation of peroxisomal proteins. In addition, since Pex5p and Pex18p are both polyubiquitinated at the N-terminal region by the same ubiquitination machinery and the fact that Cdc48p is not required for Pex18p turnover might be a point of divergence in regulation of Pex5p and Pex18p.

The current data lead to a model of peroxisomal protein quality control in which a block in Pex5p recycling leads to the accumulation of the receptor at the peroxisomal membrane. The receptor is polyubiquitinated by Ubc4p or Ubc5p (E2 enzymes) and the peroxisomal E3 ligases-complex composed out of Pex2p, Pex10p and Pex12p. In the following, Cdc48p cooperates with the heterodimeric Ufd1/Npl4 cofactor to pull misfolded, polyubiquitinated receptor out of the peroxisomal membrane for its subsequent degradation.

## Methods

### Yeast strains and culture conditions

The *S. cerevisiae* wild type strain UTL-7A (MATa, ura3-52, trp1, leu2-3/112) (Erdmann, Veenhuis et al., 1989) was used as the isogenic source for the generation of all deletion and tagged strains that are listed in table 1. Deletion and genomic integration of the GAL1-promotor 5’-of indicated coding regions were constructed by the ‘short flanking homology’ method (Güldener, Heck et al., 1996). Yeast media have been described previously (Erdmann et al., 1989). To knockdown the essential genes *CDC48*, *NPL4* and *UFD1*, galactose was used instead of dextrose to cultivate their corresponding GAL1 integrated cells.

**Table I.**
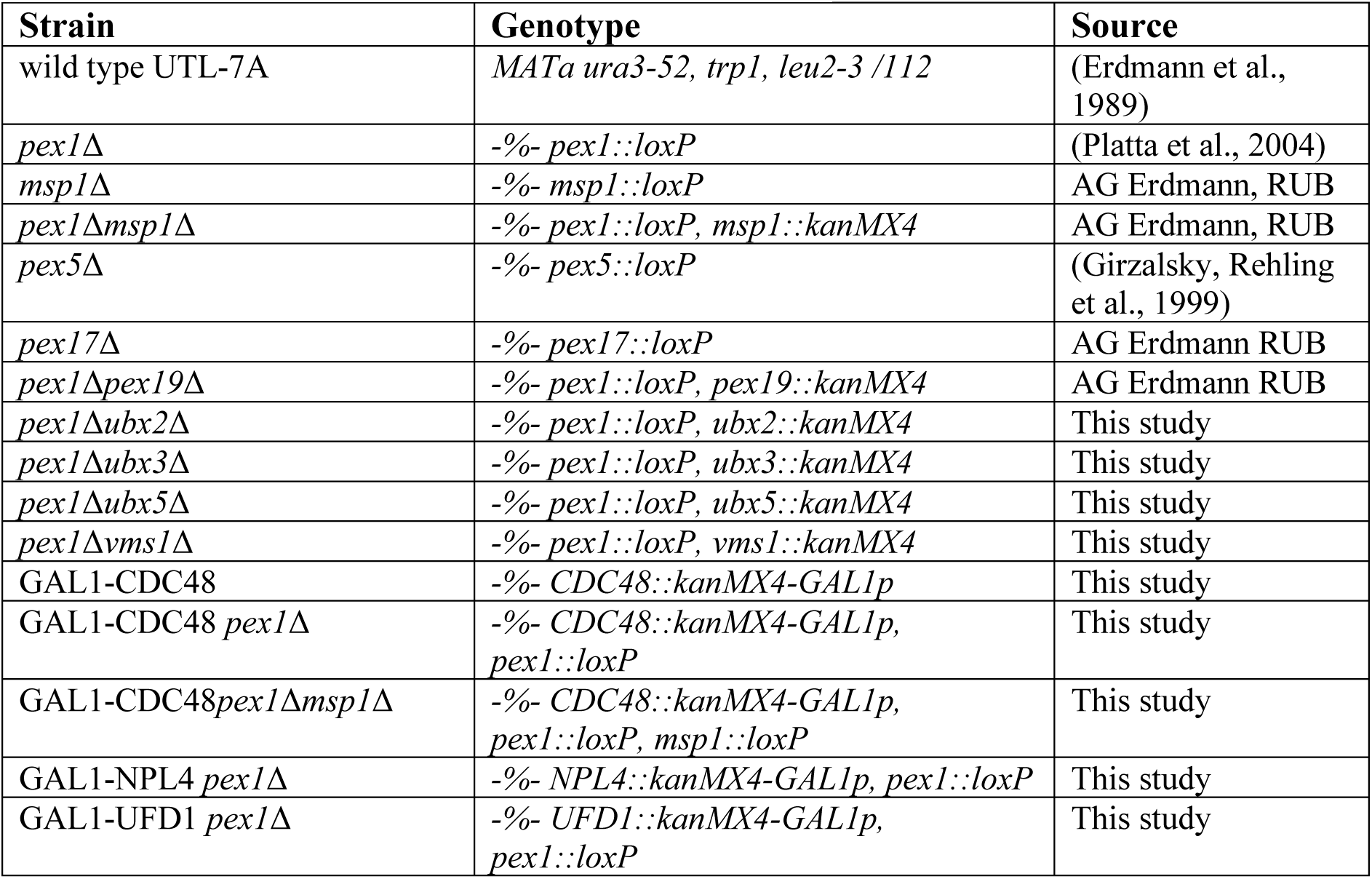
Yeast strains used in this study.

### Plasmids and cloning strategies

Plasmids and oligonucleotides used in this study are listed in tables 2 and 3, respectively. The gene encoding *S. cerevisiae* CDC48 was amplified from the yeast genomic DNA by PCR using the primer pair RE7357/RE7358 and cloned into the pRS416 vector at the BamHI and HindIII restriction sites resulting in the pRS416-CDC48 plasmid. The GAL1-promoter and the kanamycin resistance cassette were amplified by PCR from plasmid pUG6-GAL1p using corresponding primers and integrated upstream of CDC48, NPL4 or UFD1. Correct genomic integration was verified by PCR using the corresponding control primers.

**Table II.** Plasmids.

| Name | Description | Source |
| --- | --- | --- |
| pUG6 | Vector for the amplification of loxP-KanMX-loxP deletion cassette | (Güldener et al., 1996) |
| pUG6-GAL1 Promoter | KanMX-GAL1p cassette for genomic integration of GAL1p in front of Cdc48p of *S. cerevisiae* | AG Erdmann RUB |
| pRS416 | Empty vector with URA marker for generating CDC48 insert | (Sikorski & Hieter, 1989) |
| pRS416-CDC48 | Plasmid for expression of CDC48 in *S. cerevisiae* | This study |

**Table III.**
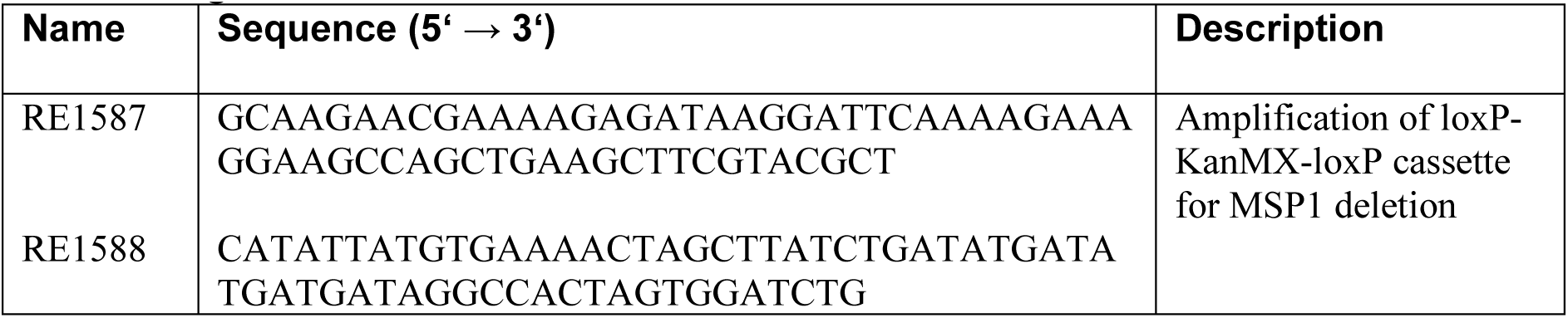
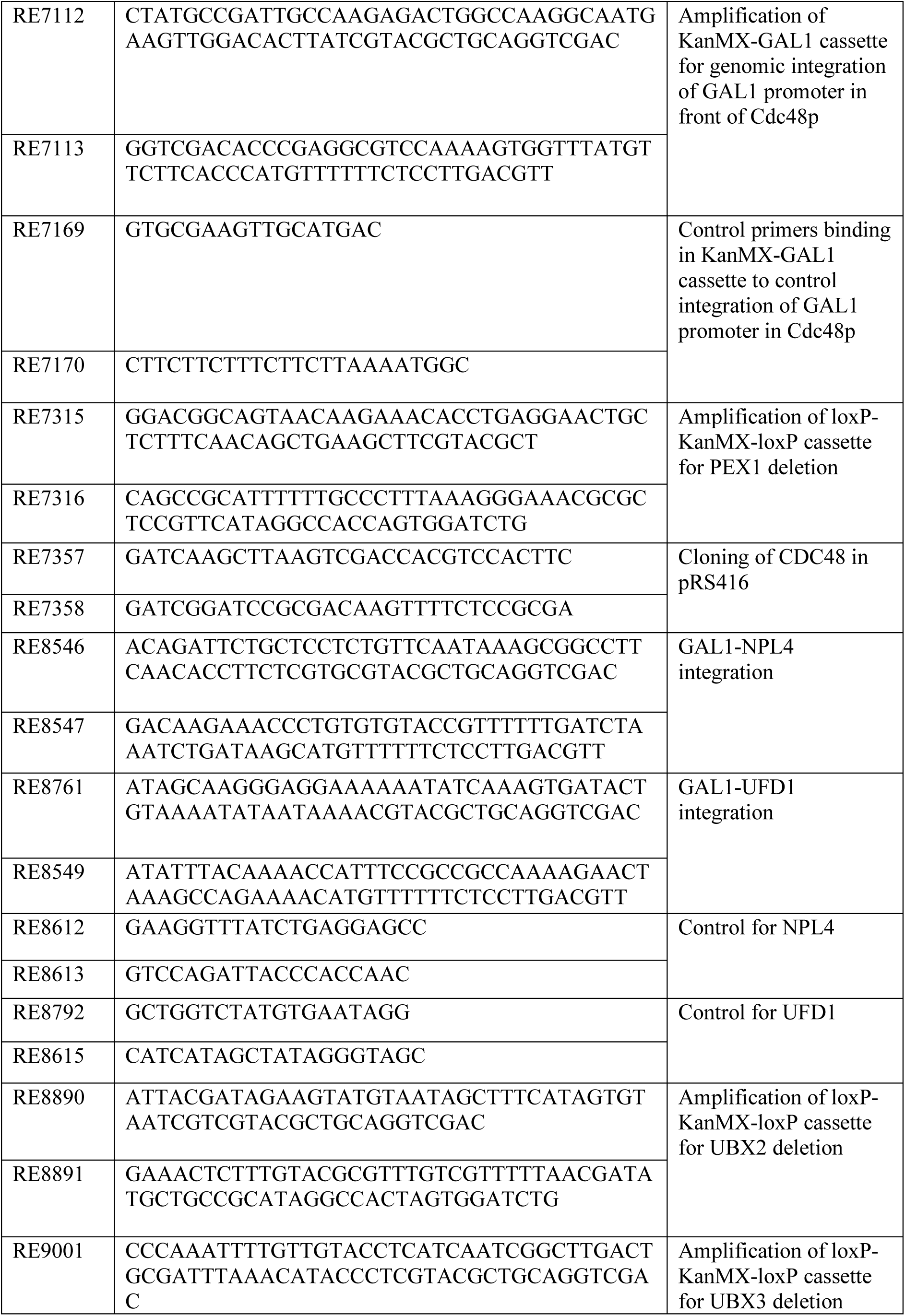
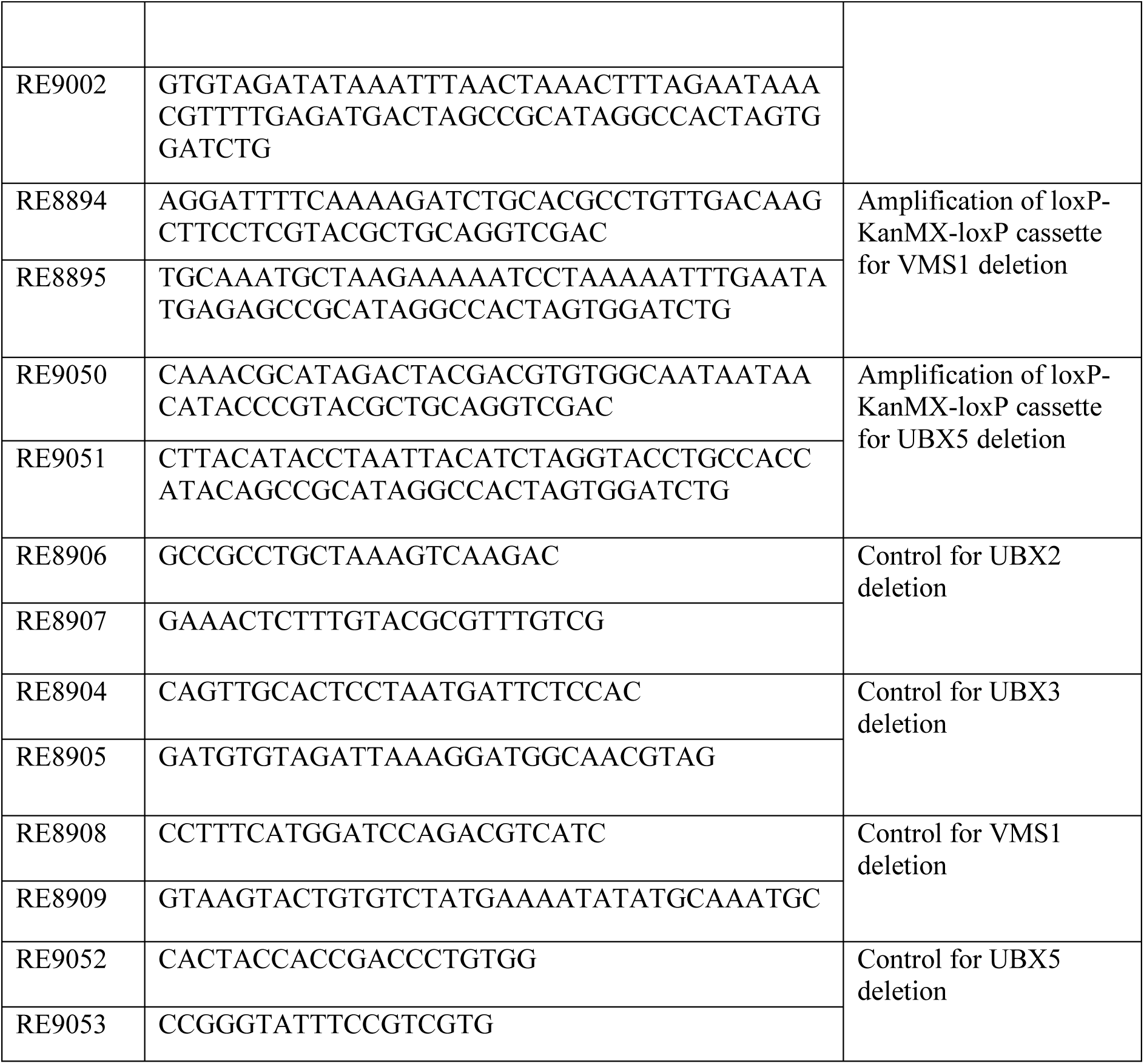
Oligonucleotides used in these studies.

### Yeast total cell extract

Yeast cells were grown at 30°C in YNB (0.5 % (w/v) ammonium sulfate, 0.17 % (w/v) YNB (Yeast Nitrogen Base, 0.1 % (w/v) yeast extract) media with an appropriate carbon source (0.3% glucose, 0.3% galactose or 0.1 % oleate). Cells were harvested, and the total protein sample was generated using a modified protocol from (Yaffe & Schatz, 1984). For this purpose, 300 µl samples of cells were prepared with an OD_600_ between 2.0 - 3.0, 15µl of potassium phosphate buffer (pH = 7.4), and 100 µl 50% (w/v) TCA were added. The samples were frozen at −80°C for at least 30 min, thawed, and centrifuged for 15 min at 13,000 rpm at 4°C. The pellet was washed with ice-cold 80% acetone and centrifuged for 10 min at 13,000 rpm, 4°C. The supernatant was removed, and the pellet dried at room temperature. The dried pellet was resuspended in 80μl of 1% (w/v) SDS/0.1M NaOH and 20μl of 5x SDS-PAGE loading buffer (5% (w/v) 2-mercaptoethanol, 15% (v/v) glycerol and 0.01% (w/v) Bromophenol Blue).

### Cycloheximid chase (CHX)

*S. cerevisiae* cells were precultured in 0.3 % glucose media and shifted to the main culture containing 0.1 % glucose and 0.1 % oleate or 0.1 % galactose and 0.1 % oleate (only for induction of Cdc48p). Samples were taken from cells grown in a culture medium to exponential or stationary growth phase (0 h). Subsequently, cycloheximide (0.5 mg/ml) or DMSO (Ctrl) was added to the culture to block protein biosynthesis. Cultures were cultivated at 30°C, and samples were taken at the indicated periods. Total cell extracts were prepared as described above and analyzed via SDS-PAGE and immunoblotting.

### Immunodetection

Protein immunodetection analyses were performed according to standard protocols (Harlow & Lane, 1988). The nitrocellulose membranes after blotting were incubated with polyclonal rabbit antisera raised against Pex5p (Albertini et al., 1997), Pex18p (Grunau, Schliebs et al., 2009), Pex11p (Erdmann & Blobel, 1995), Cdc48p (Mårtensson, Priesnitz et al., 2019), Fox3p (Erdmann & Kunau, 1994) and Porin (Kerssen, Hambruch et al., 2006). Primary antibodies were probed using the IRDye 800CW goat anti-rabbit IgG or IRDye 680RD goat anti-rabbit secondary antibody with the Odyssey® infrared imaging system (LI-COR Biosciences)

### Quantification and statistical analyses

The densitometric determination of band intensities of immunoblots was carried out with Image Studio Lite software (Ver. 5.2). The value obtained for each band was normalized with the loading control (porin) value. The standard deviations of three independent experiments were calculated using Microsoft Excel.

## Acknowledgements

The work was supported by the Deutsche Forschungsgemeinschaft, grant ER178/17-1 to RE.

## Supplementary Figures

**Supplementary Figure 1:**
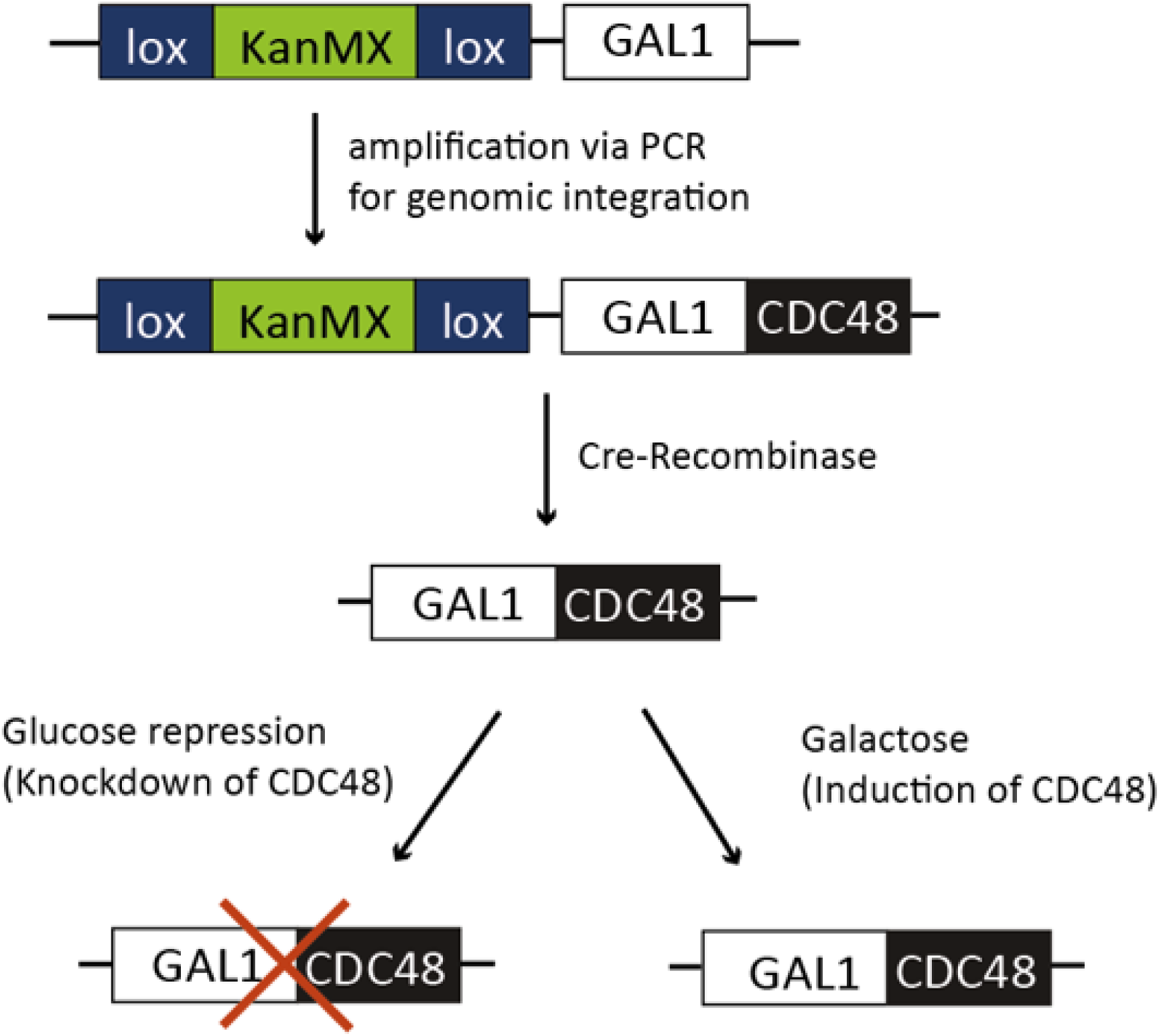
Schematic representation of the genomic integration of the GAL1 promoter upstream of CDC48. GAL1 promoter was integrated upstream of CDC48 to create CDC48 conditional knockout strains (cdc48-ck). Cdc48p-expression is switched off on glucose medium, while galactose induced Cdc48p expression.

**Supplementary Figure 2:**
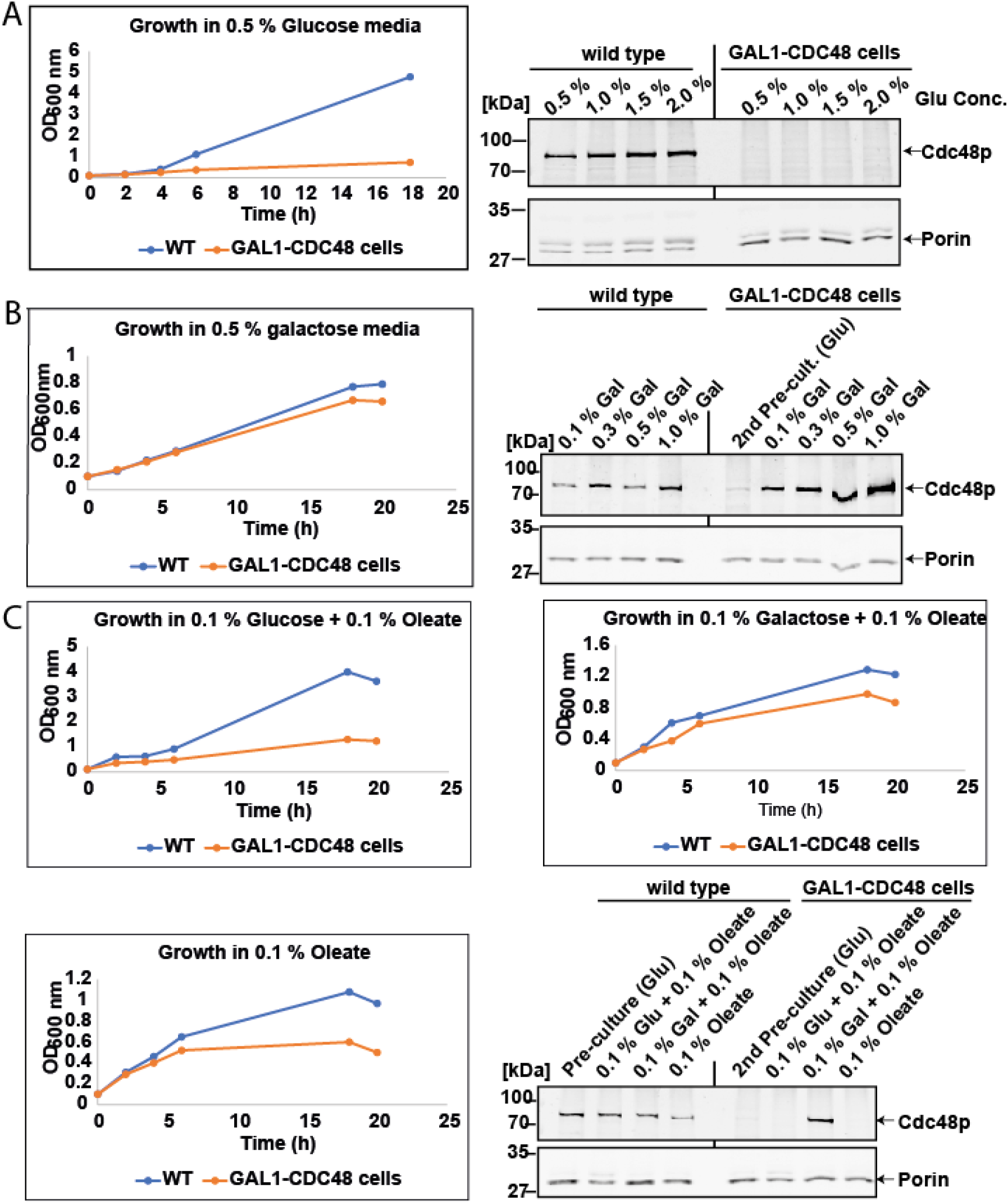
Characterization of GAL1-CDC48 integrated strains. Wildtype and GAL1-CDC48 cells were cultivated in media containing the indicated glucose concentration. (A) Growth curve with analysis of Cdc48p expression in glucose media (B) Growth curve with analysis of Cdc48p expression in galactose media (C) Growth curve with analysis of Cdc48p expression in media containing either glucose, or galactose and/or oleate galactose

**Supplementary Figure 3:**
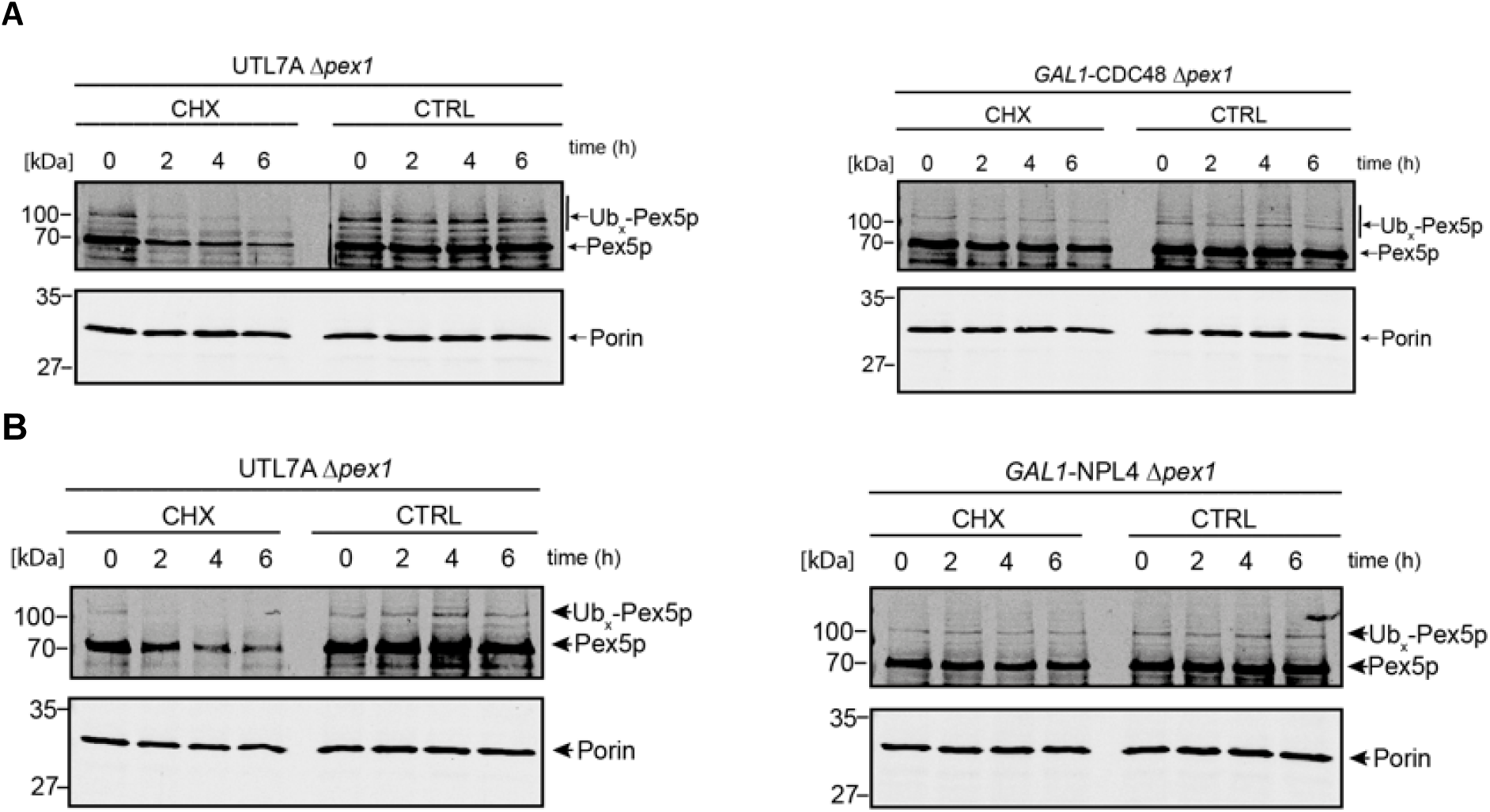
Accumulation of polyubiquitinated Pex5p. Mutant cells of *pex1Δ,* GAL1-CDC48 *pex1Δ* and GAL1-NPL4 *pex1Δ* cells were cultivated in media containing 0.1 % glucose and 0.1 % oleate to deplete Cdc48p (A) or Npl4p (B). All strains were cultivated to the exponential phase (OD600nm = 1-1.5) and then treated with DMSO (Ctrl) or 0.5 mg/ml cycloheximide (CHX) as indicated. Cell lysates were prepared by TCA precipitation at the indicated times, and proteins were analysed by immunoblotting using the indicated antibodies. The data show that the depletion of both Cdc48p and Npl4p results in a stabilization of Pex5p and the appearance of higher-molecular weight species of Pex5p that are known to represent poly-ubiquitinated forms.

**Supplementary Figure 4:**
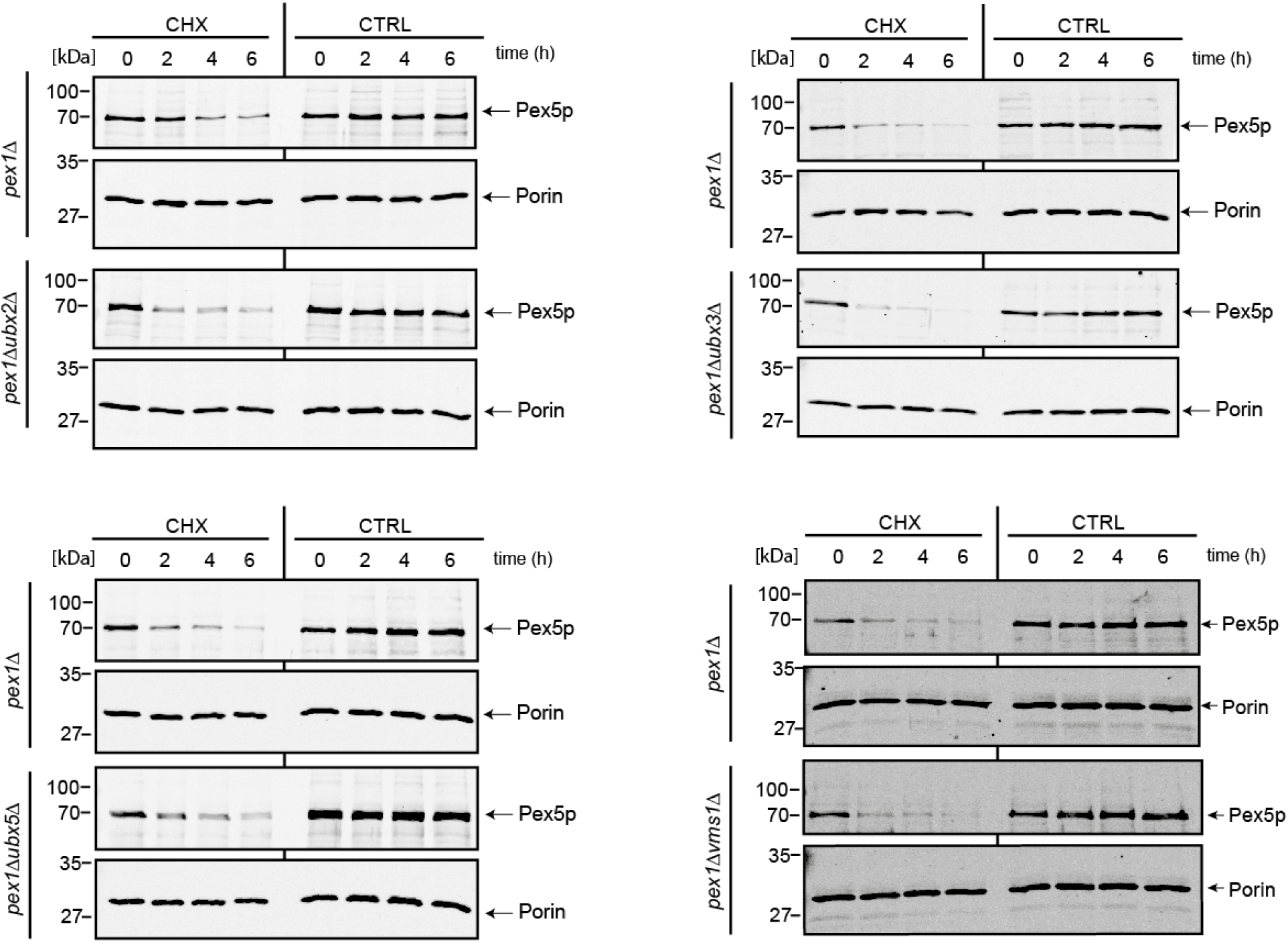
Cdc48p co-factors Ubx2, Ubx3, Ubx5 and Vms1 are not required for Pex5p turnover. *pex1Δ*, *pex1Δubx2Δ, pex1Δubx3Δ, pex1Δubx5Δ* and *pex1Δvms1Δ* cells were cultivated in media containing 0.1 % glucose and 0.1 % oleate to the exponential phase (OD600nm = 1-1.5) and then treated with DMSO (Ctrl) or 0.5 mg/ml cycloheximide (CHX). Cell lysates were prepared by TCA precipitation at the indicated times, and proteins were analysed by immunoblotting using the indicated antibodies.

